# Unheeded SARS-CoV-2 proteins? A deep look into negative-sense RNA

**DOI:** 10.1101/2020.11.27.400788

**Authors:** Martin Bartas, Adriana Volná, Christopher A. Beaudoin, Ebbe Toftgaard Poulsen, Jiří Červeň, Václav Brázda, Vladimír Špunda, Tom L. Blundell, Petr Pečinka

## Abstract

SARS-CoV-2 is a novel positive-sense single-stranded RNA virus from the *Coronaviridae* family (genus *Betacoronavirus*), which has been established as causing the COVID-19 pandemic. The genome of SARS-CoV-2 is one of the largest among known RNA viruses, comprising of at least 26 known protein-coding loci. Studies thus far have outlined the coding capacity of the positive-sense strand of the SARS-CoV-2 genome, which can be used directly for protein translation. However, it has been recently shown that transcribed negative-sense viral RNA intermediates that arise during viral genome replication from positive-sense viruses can also code for proteins. No studies have yet explored the potential for negative-sense SARS-CoV-2 RNA intermediates to contain protein coding-loci. Thus, using sequence and structure-based bioinformatics methodologies, we have investigated the presence and validity of putative negative-sense ORFs (nsORFs) in the SARS-CoV-2 genome. Nine nsORFs were discovered to contain strong eukaryotic translation initiation signals and high codon adaptability scores, and several of the nsORFs were predicted to interact with RNA-binding proteins. Evolutionary conservation analyses indicated that some of the nsORFs are deeply conserved among related coronaviruses. Three-dimensional protein modelling revealed the presence of higher order folding among all putative SARS-CoV-2 nsORFs, and subsequent structural mimicry analyses suggest similarity of the nsORFs to DNA/RNA-binding proteins and proteins involved in immune signaling pathways. Altogether, these results suggest the potential existence of still undescribed SARS-CoV-2 proteins, which may play an important role in the viral lifecycle and COVID-19 pathogenesis.

## Introduction

The Severe Acute Respiratory Syndrome Coronavirus – 2 (SARS-CoV-2) has been intensively studied worldwide for its role as the causative agent of the COVID-19 pandemic (Hu et al. 2020; Wu et al. 2020; Zheng 2020). Coronaviruses, such as SARS-CoV-2, are single-stranded positive-sense RNA viruses and have the largest genomes among RNA viruses – usually around 30 kb. It has been established that the SARS-CoV-2 genome codes for at least 26 proteins: 16 non-structural proteins (NSP1-16), 4 structural proteins (surface glycoprotein, membrane glycoprotein, envelope protein, nucleocapsid phosphoprotein), and 6 – 9 accessory factors (designated as open reading frames, ORFs) (Gordon et al. 2020; Jiang et al. 2020; Zhang et al. 2020; Finkel et al. 2021). Additionally, several overlapping genes and new ORFs, which may code for new proteins, have been discovered in the accessory region in recent months as well, suggesting that the SARS-CoV-2 proteome is not completely resolved (Nelson et al. 2020)

Positive-sense RNA viruses, such as SARS-CoV-2, have been thought to encode proteins solely on the positive strand. However, Dinan *et al.* recently demonstrated that negative-sense viral RNA strand intermediates arising during replication of viral positive-sense RNA genomes also have protein-coding potential (Dinan et al. 2020). Previous ribo-seq analysis of an infection model with the murine coronavirus (mouse hepatitis virus, strain A59) revealed that negative-sense RNA was found at significantly lower levels than the positive-sense and that translation on the negative strand was uncertain (Irigoyen et al. 2016). Although studies have quantified the amount of SARS-CoV-2 negative-sense RNA in host cells, which is present at approximately 10 to 100 times lower than positive-sense RNA, no studies to-date have described the potential for coding sequences on the negative strand of the SARS-CoV-2 genome (Alexandersen et al. 2020).

Herein, a combination of complementary sequence and structure-based bioinformatics approaches was used to elucidate the presence of protein-coding negative-sense ORFs (nsORFS) in the SARS-CoV-2 genome. We, first, identified and cross-examined the presence of eukaryotic translation initiation sites (Gong et al. 2014; Noderer et al. 2014) and open reading frames on the SARS-CoV-2 negative-sense genome using four distinct tools. The predicted nsORFs were then subjected to codon bias analysis, transcription factor binding site analysis, sequence and domain-based homology searches, proteomics meta-searches, ribosome profiling analysis, and 3D structure prediction and characterization to understand their potential validity and functionality. In summary, we discovered nine putative protein-coding regions on the negative-sense SARS-CoV-2 RNA that exhibited codon biases consistent with the human genome and were predicted to contain higher-order 3D structural folding. We extended our reach to check for nsORFs in phylogenetically related coronavirus genomes and discovered that the presence of protein-coding regions on negative-sense coronavirus RNA may be evolutionarily conserved and widespread. Proteomics and Ribo-Seq analyses were unable to detect whether these nsORFs are translated during infection; however, because of the low amount of negative-sense RNA, detection of translation may require more focused and in-depth experimentation. Our analyses propose novel SARS-CoV-2 ORFs that may play a role during infection of host cells.

## Results and Discussion

### Novel ORFs with Kozak consensus sequences detected on SARS-CoV-2 negative-sense strand

The detection of translation initiation sequences in viral genomes for the prediction and characterization of potential protein-coding sequences has been described for several viral pathogens (Nair et al. 2016; Goldberg et al. 2019; La Bella et al. 2020; Tan et al. 2021). In order to detect potential coding sequences on the SARS-CoV-2 negative-sense genome, we used two web servers, TISrover (Zuallaert et al. 2018) and ATGpr (Salamov et al. 1998), that detect eukaryotic ribosome translation initiation sites (TIS) based on machine learning algorithms and two web servers, NCBI ORFfinder (https://www.ncbi.nlm.nih.gov/orffinder/) and StarORF (http://star.mit.edu/orf/index.html), that look for ORFs based on six-frame translation of nucleotide sequences. The TIS detection tools search for eukaryotic translation start signals, such as the Kozak sequence (A/GXXATGG), which have been recorded to significantly affect gene expression (Acevedo et al. 2018; Jaafar and Kieft 2019). The TIS detection tools provide confidence scores from 0-1 that can be used to discern the probability of the predicted start site. The SARS-CoV-2 positive strand and its recorded gene start sites were analyzed in parallel using the TIS detection tools as a control measure and to set threshold values for the TIS detection tools (Nelson et al. 2020; Wu et al. 2020). The TIS detection tools correlated well with the positive strand gene start sites (Supplementary Table S1). The first 8 start sites found using ATGpr corresponded to the M, ORF9b, N, pp1a, orf1ab, ORF8, truncated version of N, and S genes, while also detecting the ORF3a, ORF7a, and ORF9c genes above the 0.1 score. TISrover presented lower sensitivity but still detected the M, N, ORF7b, orf1ab, pp1a, ORF6, ORF8, and ORF3a start sites at above a 0.1 score. We, thus, set a threshold value of 0.1 for both ATGpr and TISrover for detection of putative TIS sites on the negative strand. A value of 0.1 has also been established as a threshold using ATGpr in other human and viral TIS detection studies (Monjaret et al. 2014; Hickman et al. 2018). Three criteria were established for selection of potential ORFs: the sequences must be 1) found using all four tools or 2) found in both TIS detection tools above the 0.1 threshold, and 3) sequence length must be above 40 amino acids. After filtering based on the criteria, 9 sequences were selected to be potentially protein-coding on the negative strand of SARS-CoV-2. Each of the nine had a strong Kozak signal, a stop codon, and ranged from 132-300 nt (Table 1).

**Table 1.** Sequence position and analysis of nsORFs

| nsORF | Frame | Identity to Kozak rule (A/GXXATGG) | Start (bp) | Finish (bp) | Length (nuc/aa) | ATGpr score | TISrover score | CAI |
| --- | --- | --- | --- | --- | --- | --- | --- | --- |
| nsORF1 | 2 | AXXATGc | 562 | 694 | 132 / 44 | 0.1 | 0.861 | 0.717 |
| nsORF2 | 2 | tXXATGt | 2899 | 3097 | 198 / 66 | 0.06 | 0.008 | 0.712 |
| nsORF3 | 3 | cXXATGa | 5792 | 5975 | 183 / 61 | 0.09 | 0.028 | 0.693 |
| nsORF4 | 2 | tXXATGa | 6466 | 6703 | 237 / 79 | 0.16 | 0.102 | 0.806 |
| nsORF5 | 1 | AXXATGa | 8865 | 9057 | 192 / 64 | 0.09 | 0.194 | 0.654 |
| nsORF6 | 1 | GXXATGt | 10047 | 10188 | 141 / 47 | 0.11 | 0.909 | 0.739 |
| nsORF7 | 3 | AXXATGt | 23414 | 23714 | 300 / 100 | 0.22 | 0.015 | 0.682 |
| nsORF8 | 1 | cXXATGa | 29211 | 29385 | 174 / 58 | 0.14 | 0.232 | 0.705 |
| nsORF9 | 2 | AXXATGG | 29236 | 29479 | 243 / 81 | 0.47 | 0.889 | 0.776 |

The predicted negative-sense ORFs (nsORFs) were numbered in order of their appearance in the 5’→3’ direction of the negative-sense RNA (nsORF1-9). The putative nsORFs are found distributed throughout the negative-sense strand and two, nsORF8 and nsORF9, are overlapping on different reading frames. Based on the 5’→3’ directionality of the positive strand, 5 of the nsORFs are found within the orf1ab region, and the remaining 4 are found amongst the structural and accessory protein genomic regions (Figure 1). Amino acid sequence-based similarity detection tools (Pfam, SMART, and others) were unable to detect homologous genes for all predicted nsORFs.

**Figure 1.**
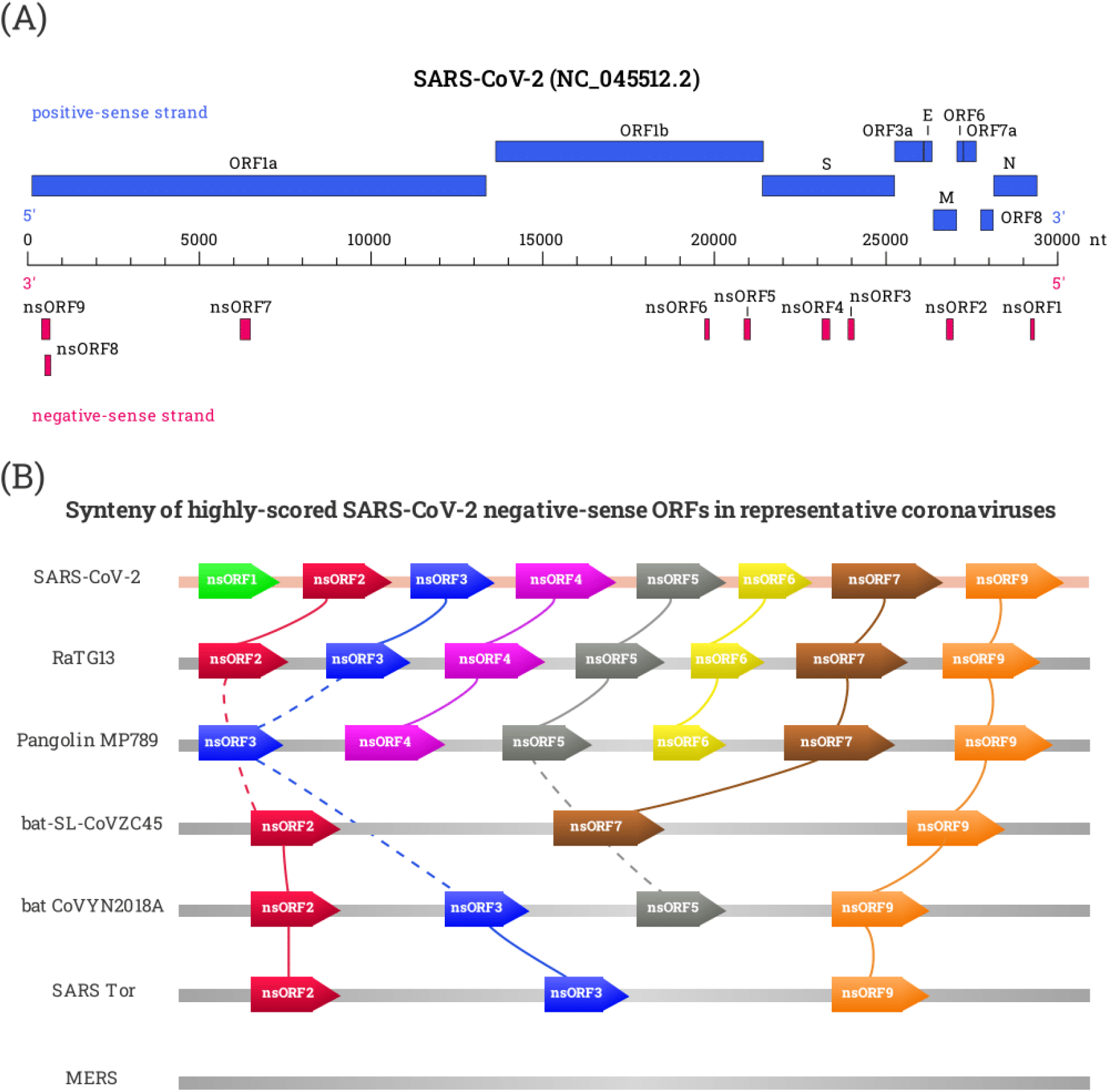
Localization and synteny of nsORFs in SARS-CoV-2 and related coronaviruses. (A) Localization of all identified nsORFs within the SARS-CoV-2 genome. (B)Synteny of SARS-CoV-2 nsORFs in representative species of SARS-like coronaviruses. At least two SARS-CoV-2 nsORFs (nsORF2 and nsORF9) are more or less conserved in most of inspected SARS-CoV-2-related coronaviruses, including old SARS-CoV Tor (2003). In MERS-CoV and human-CoV-OC43, none of SARS-CoV-2 homologous nsORFs was found. The synteny plot was constructed using SimpleSynteny web server (Veltri et al. 2016) and redrawn to this schematic figure.

To explore whether predicted nsORFs contain binding motifs for human proteins, further bioinformatic analysis utilizing Beam RNA Interaction motif search tool (Guarracino et al. 2021) was conducted. This RNA interaction motif analysis revealed many interesting hits: nsORF9 (and its overlapping nsORF8) contains a motif that is significantly similar to the PUM1 binding sequence (p-value = 0.012). The PUM1 protein has been reported to play a role in cytoplasmic sensing of viral infection (Narita et al. 2014). nsORF7 contains a motif that is significantly similar to the UPF1 binding sequence (p-value = 0.019); this protein is also called the “Regulator of nonsense transcripts 1” and plays a vital role in host-virus interaction (May and Simon 2021). nsORF6 contains a sequence motif that is significantly similar to the MOV10 binding sequence (p-value = 0.0083), and MOV10 has been identified to exhibit antiviral activity against dengue virus (which is also a positive-sense ssRNA virus) (Balinsky et al. 2017). Interestingly, MOV10 is a “5′ to 3′ RNA helicase contributing to UPF1 mRNA target degradation by translocation along 3′ UTRs” (Gregersen et al. 2014: 10). nsORF5 contains a motif that is significantly similar to the ATP-dependent RNA helicase SUPV3L1 binding sequence (p-value = 0.012); because this protein is considered to be mitochondrial (Szczesny et al. 2010), the interaction with SARS-CoV-2 RNA seems to be unlikely. nsORF4 contains a motif that is significantly similar to the heterogeneous nuclear ribonucleoprotein L (hnRNP L) binding sequence (p-value = 0.02). Notably, it was previously reported that hnRNP L interacts with hepatitis C virus (positive-sense ssRNA virus) 5′-terminal untranslated RNA and promotes efficient replication (Li et al. 2014). nsORF3 contains a motif that is significantly similar to the U2AF5 binding sequence (p-value = 0.0091) and also a motif that is significantly similar to the hnRNP L binding sequence (p-value = 0.025), as in nsORF4. nsORF2 contains a motif that is significantly similar to the GRWD1 binding sequence (p-value = 0.00022), but the functions of this protein are still largely unknown. nsORF1 contains motifs that are significantly similar to nuclear cap-binding protein subunit 3 (NCBP3) binding sequence (p-value = 0.034) and U2AF2 binding sequence (p-value = 0.04). NCBP3 associates with NCBP1/CBP80 to form an alternative cap-binding complex which plays a key role in mRNA export; it is also known that the alternative cap-binding complex is important in cellular stress conditions such as virus infections and the NCBP3 activity inhibit virus growth (Gebhardt et al. 2015).

Codon-usage similarity between viral and host genomes has been shown as an indicator for adaptation to the host, since optimized use of the available endogenous amino acids allows more efficient translation of viral genes (Bahir et al. 2009; Jitobaom et al. 2020). The codon-usage of the canonical SARS-CoV-2 genes has been determined to correlate well with the human, bat, and pangolin amino acid pools (Gu et al. 2020; Roy et al. 2021). Using the codon adaptability index (CAI), which has been shown as an accurate predictor for gene expression levels, studies have shown that the CAI values of the positive strand average around 0.7 (with 1 being the best score) (Sharp and Li 1987; Dilucca et al. 2020; Li et al. 2020). Thus, to better understand the expression efficiency of the putative nsORFs and their codon usage similarity to the human amino acid pool, codon usage tables were created and analyzed using COUSIN and the CAI values for the nsORFS were calculated using the CAIcal web server (Puigbò et al. 2008; Bourret et al. 2019). Comparison of the relative frequencies of codons used by the positive and negative-sense genomes in relation to the human genome revealed a high correlation between preferred codons. As shown in Table 1, average CAI values for the positive strand were 0.68 and ranged from 0.606-0.726, while the average for negative strand ORFs was 0.72 and ranged from 0.654-0.806. Notably, nsORF4, nsORF6, and nsORF9 reported higher CAI values (0.806, 0.739, and 0.776 respectively) than the maximum reported CAI for the positive strand genes (N protein: 0.726). The high congruence between the CAI values of the negative and positive-sense ORFs to the human amino acid pool lends further evidence for potential expression of these genes.

In order to detect whether the nsORFs are translated in human cells, we performed 1) proteomics meta-searches of the mass spectrometry data from two other studies involving SARS-CoV-2 infection of human primary alveolar macrophages (Dalskov et al. 2020) and Vero E6 cells (Grenga et al. 2020); and 2) an analysis of ribosomal profiling data from Finkel et al. (Finkel et al. 2021). Unfortunately, the signals were too weak in both cases to confirm translation. Studies have shown that lowly expressed proteins, such as the E protein (only 20 copies per virion (Bar-On et al. 2020)), may not be discovered using proteomics techniques (Gouveia et al. 2020; Renuse et al. 2021). Additionally, negative-sense RNA has been to present at 10 to 100 times lower than the amount of positive-sense RNA (Alexandersen et al. 2020). The Ribo-Seq data reflected this pattern, as the gene transcript mapping failed to attain a threshold level of genome coverage (Nelson et al. 2020). Thus, more focused or high-depth Ribo-Seq profiling or proteomics may better resolve the *in vivo* presence of these proteins.

### Evolutionarily conservation of nsORFs

Evolutionarily conservation of ORFs has been considered as supporting evidence for protein expression (Shi et al. 2012). Thus, we used SimpleSynteny tool (Veltri et al. 2016) to investigate nsORF synteny among coronaviruses. We have found, that in the closest relative, RaTG13 coronavirus genome, nearly all nsORFs (except of nsORF1) are conserved and not truncated by stop codons (Figure 1). In more distant coronaviruses (e.g. Bat SARS-like coronavirus isolate bat-SL-CoVZC45, coronavirus BtRs-BetaCoV/YN2018A, and SARS coronavirus Tor2), three to four SARS-CoV-2 nsORFs still have their homologs (Figure 1). Interestingly, nsORF3 and nsORF5 were truncated in Bat SARS-like coronavirus isolate bat-SL-CoVZC45, but preserved in evolutionarily more distant coronavirus BetaCoV/YN2018A. nsORF2 and nsORF9 were conserved in all inspected viral strains of SARS-related coronaviruses, and these are SARS-CoV-2 nsORFs identified by all four approaches – TISrover, ATGpr, NCBI ORFfinder, and StarORF (Fuchs et al. 2001). In MERS-CoV, human-CoV-OC43, and more distant members of *Coronaviridae* family, no homologous SARS-CoV-2 nsORFs were found.

### ORFs predicted to contain higher order folding: modelling, characterization, and comparison

To gain more insight into the potential functionality of these genes, despite the uncertainty of their translation, we predicted the 3D structure of each nsORF, characterized the predicted structures, and performed structural comparisons with all 3D experimentally-resolved proteins. As no templates were available for homology modelling, *ab initio* structural modelling with trRosetta (Yang et al. 2020) was used in combination with secondary structure prediction and structural refinement with RaptorX (Källberg et al. 2012) and MODELLER (Sali and Blundell 1993; Webb and Sali 2017), respectively. All nsORFs were predicted to have higher order folding (Figure 2). Potential transmembrane region analysis using TMHMM and the OPM database revealed that only nsORF7 was predicted to contain a transmembrane domain (Figure 2A) (Krogh et al. 2001; Lomize et al. 2012). To study the effect of major post-translational modifications, N- and O-linked glycosylation motifs were detected using NetNGlyc (Gupta and Brunak 2001) and NetOGlyc (Steentoft et al. 2013) and modelled using the CHARM-GUI Glycan Reader and Modeler (Park et al. 2019). nsORF9 and nsORF6 were predicted to have one and two N-linked glycosylation sites, respectively, and nsORF5 and nsORF3 were predicted to contain 10 and 4 O-linked glycosylation sites, respectively (Figure 2B). Heavy glycosylation may imply potential roles in inflammatory processes as secreted signaling proteins (Reily et al. 2019). Isoelectric points, predicted by ExPASy (Gasteiger et al. 2005), were found at an average 9.07, which is reflected by the higher presence of basic residues, as shown in Figure 2C. The presence of positively charged residues may have implications in viral or host nucleic acid binding (Komazin-Meredith et al. 2008; Requião et al. 2020). Overall, the nsORFs were found to contain higher order folding and several structural characteristics of interest.

**Figure 2.**
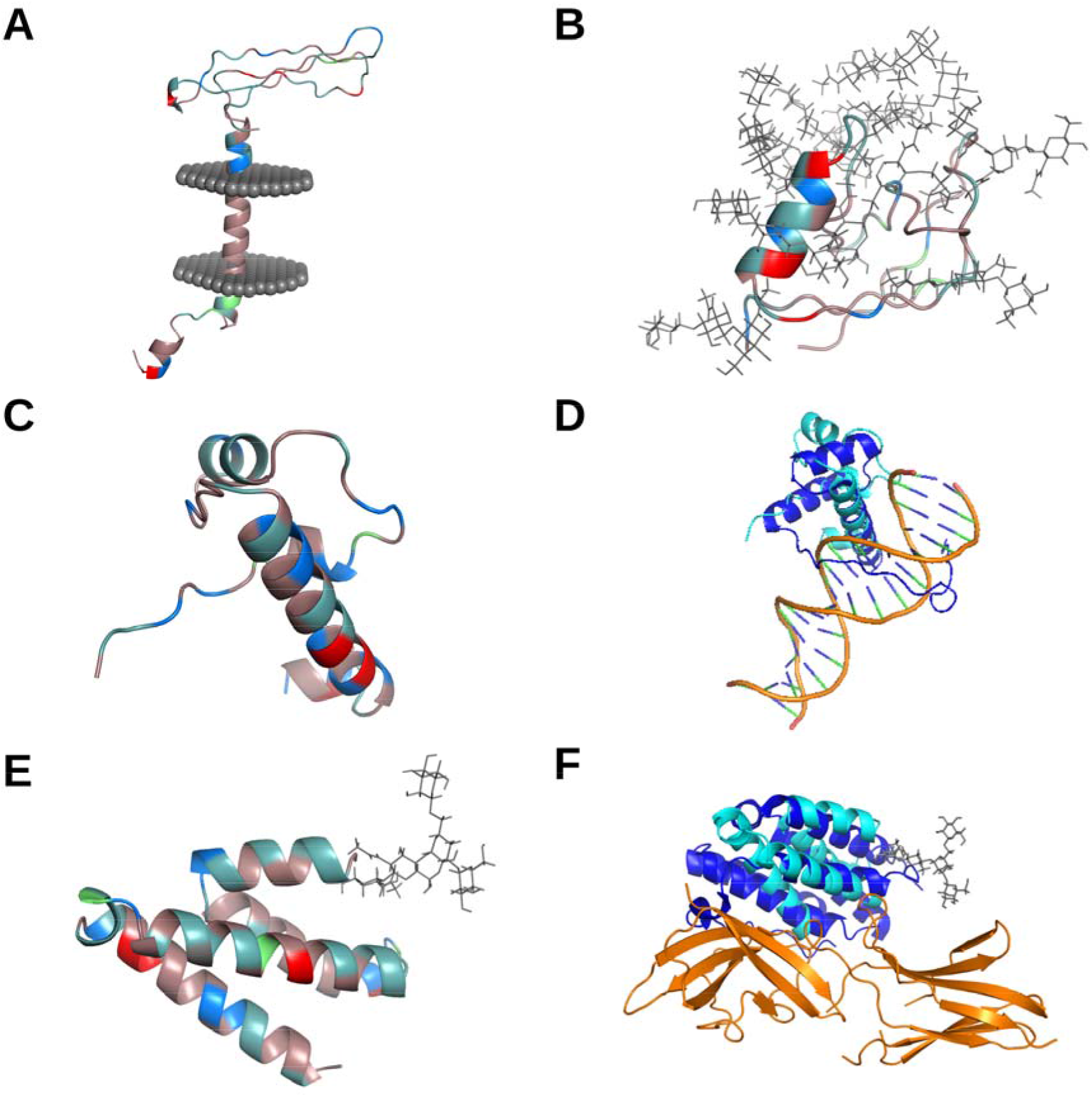
Structural characterization and similarity comparisons of nsORFs Residues of putative nsORFs in A (nsORF7), B (nsORF5), C (nsORF1), and E (nsORF9) are depicted with amino acid coloration: red for acidic (D,E), blue for basic (H,R,K), light teal for polar non-charged (S,N,T,Q), dirty violet for hydrophobic (A,V,I,L,M,F,W,P,G,Y), and lime green for cysteine residues. Putative transmembrane protein nsORF7 is shown with the predicted transmembrane region inside a representative cell membrane (A). Extensive O-linked glycosylation of nsORF5 is shown with grey stick configurations (B). The structural similarity of nsORF1 (C) to a homologous protein of T-cell leukemia homeobox protein 2 (which was predicted by RUPEE, but shown without DNA) bound to DNA (PDB: 3a01; both homeobox protein structures are published by (Miyazono et al. 2010) is depicted with nsORF1 in cyan, homeobox protein protein in blue, and DNA in orange (D). The predicted protein-protein interaction of nsORF9 (E) and interferon alpha/beta receptor 2 using HMI-PRED is compared to the interaction between interferon alpha/beta receptor 2 and interferon omega-1 (PDB: 3se4) with nsORF9 in cyan, interferon omega-1 in blue, and interferon alpha/beta receptor 2 in orange (F).

Structural similarity comparisons have been shown to give insight into potential protein-protein interactions, despite low sequence similarity (Drayman et al. 2013; Beaudoin et al. 2021). Using RUPEE (Ayoub and Lee 2019), the nsORFs were compared to all known protein families, and HMI-PRED (Guven-Maiorov et al. 2020) was used to infer potential host interaction partners. Structural alignments generated using RUPEE revealed that three nsORFs, 2, 4 and 9, exhibited high structural similarity to known proteins with TM-scores over 0.5 (indicating that they are in the same fold), while nsORFs 1, 3, 5 and 8 reported the lowest similarity with TM-scores under 0.4 (Zhang and Skolnick 2005). The highest returned TM-scores of the nsORFs, such as 1, 4, and 9, were predicted to be structurally similar to RNA/DNA binding proteins (T-cell leukemia homeobox protein 2, DNA-Binding Domain of mouse MafG, and RNA Binding Domain from influenza virus nonstructural protein 1, respectively), furthering evidence from the isoelectric point observations (Figure 2D) (Miyazono et al. 2010). Cell signaling factors, such as those involved in complement activation, may be mimicked by nsORF6 and nsORF8, while proteins involved in protein degradation and other ubiquitin-related processes might interact with nsORF2, nsORF3, nsORF6, and nsORF9 (Table 2). Interestingly, only nsORFs 2, 4, 7, and 9 returned potential mimicked/disrupted protein-protein interfaces by HMI-PRED (Supplementary Table S2). Diverse cellular processes were predicted to be involved in the mimicked interfaces ; for instance, nsORF9 was found structurally similar to interferon alpha/beta receptor 2 binding to interferon omega-1, which could have roles in inflammatory signaling in SARS-CoV-2 infections (Figure 2F) (Thomas et al. 2011). The higher order folding of these ORFs and similarity to known proteins provides further evidence that they may have functional roles in infection.

**Table 2.**
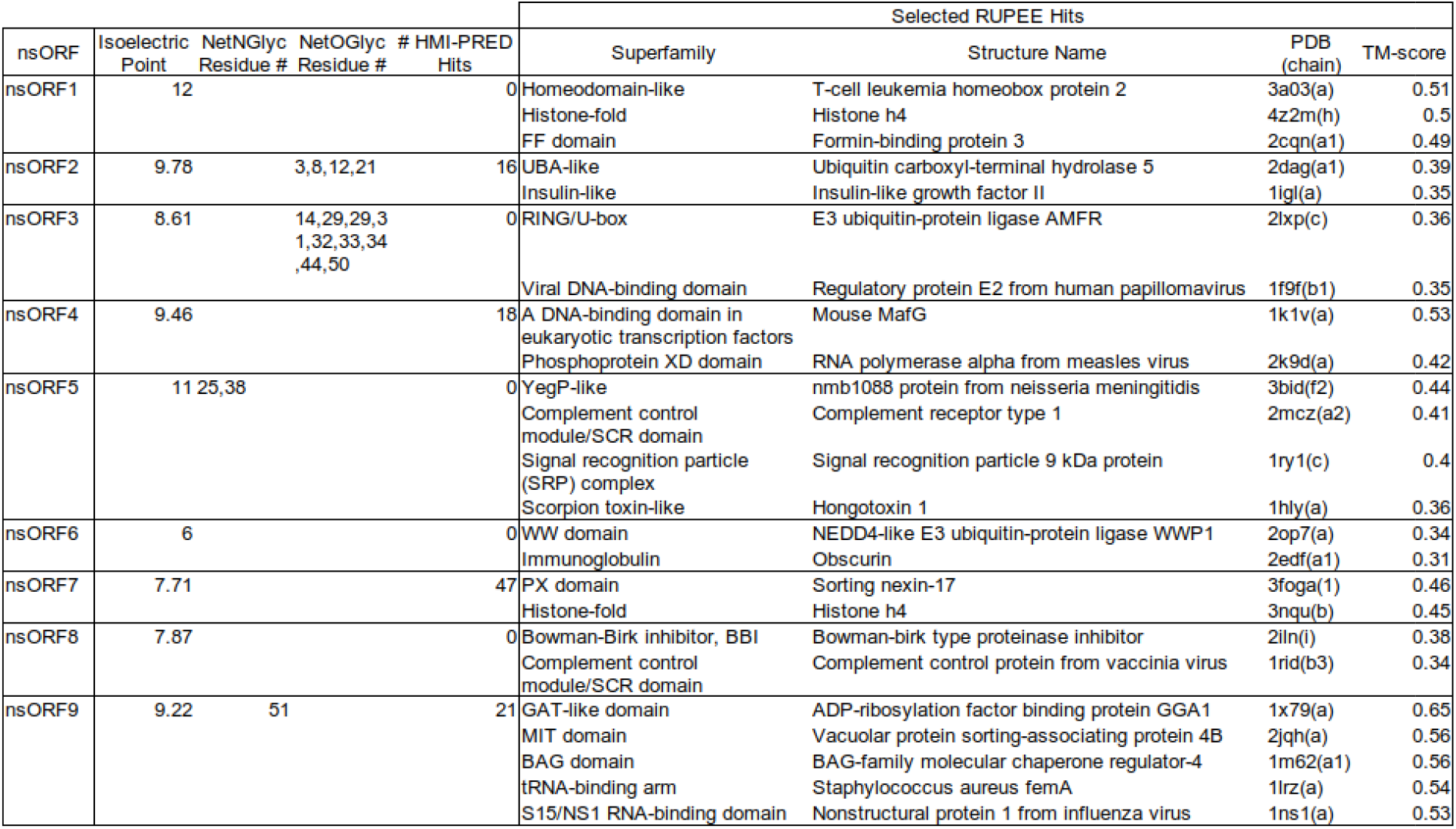
Structural characterization of nsORFs

## Conclusion

Altogether, our results suggest the existence of still undescribed SARS-CoV-2 proteins, which may play an important role in the viral lifecycle and COVID-19 pathogenesis. Nine potential nsORFs were discovered using various sequence and structure-based bioinformatics methodologies. The nsORFs were unable to be detected using publicly-available proteomics or ribosomal profiling datasets, which may reflect their low overall abundance. Interestingly, the average codon adaptability of the nsORFs was higher than that of the positive-sense SARS-CoV-2 genes, which may be a compensatory mechanism to account for low levels of negative-sense RNA as templates for translation. All nine nsORFs were predicted to have higher order folding, which was confirmed by the structural similarity to several known human and viral proteins. For example, both nsORF2 and nsORF9 were predicted to have histone-like folds. Furthermore, nsORF2 contains sorting nexin-like fold, and nsORF9 contains formin-binding-like fold. Notably, sorting nexin proteins and formin-binding proteins are known to be interaction partners, which may give more indications for their complementary roles during infection (Fuchs et al. 2001). Is it therefore possible, that some of the SARS-CoV-2 nsORFs are expressed and form protein complexes, similarly as nsp1-nsp16 on the positive SARS-CoV-2 RNA strand? We hope that this study will stimulate further research in the field of developing more specific and sensitive approaches to detect the complete SARS-CoV-2 proteome *in vitro* and *in vivo*.

## Materials and Methods

### Sequence collection and ORF detection and characterization

The SARS-CoV-2 reference genome (NC_045512.2) was selected and reverse-transcribed using Reverse complement tool (https://www.bioinformatics.org/sms/rev_comp.html) as a reference for the negative-sense strand. A combination of four tools was used to discover ORFs and Kozak sequences on the negative-sense strand: TISRover prediction tool for predicting translation initiation sites in human by convolutional neural networks (Zuallaert et al. 2018); ATGpr tool (https://atgpr.dbcls.jp/) which uses linear discriminant analysis for identifying the initiation codons (Salamov et al. 1998); NCBI ORFfinder (https://www.ncbi.nlm.nih.gov/orffinder/) to predict all potential ORFs; and StarORF (http://star.mit.edu/orf/index.html) to cross examine ORFfinder results. The RegRNA 2.0. webserver (Chang et al. 2013) and Beam RNA Interaction mOtif search tool (BRIO) (Guarracino et al. 2021) were used for the transcription factor binding site analysis. Codon usage tables were made using COUSIN (Bourret et al. 2019), and the codon adaptability index (CAI) was calculated using the CAIcal web server (Puigbò et al. 2008).

### ORFs conservation and synteny in related viral species

To inspect, whether there are proteins homologous to the SARS-CoV-2 negatively-encoded ORFs also in another important coronaviral species, we made tblastn searches (https://blast.ncbi.nlm.nih.gov/Blast.cgi) using negatively-encoded ORFs (protein sequences) as a query. The searches and further analyses were restricted to the following representative betacoronaviral (β-CoV) genomes: Bat coronavirus RaTG13 (MN996532.2), Bat SARS-like coronavirus isolate bat-SL-CoVZC45 (MG772933.1), SARS coronavirus Tor2 (NC_004718.3), Coronavirus BtRs-BetaCoV/YN2018A (MK211375.1), MERS-CoV isolate HCoVEMC2012 (NC_019843), and Human-CoV-OC43 (NC_005147.1). Synteny was analyzed and graphically depicted using the SimpleSynteny tool (https://www.dveltri.com/simplesynteny/about.html) (Veltri et al. 2016).

### Structural characterization of potential protein-coding sequences

To find possible domain homologs we used NCBI’s Conserved Domains Database (CDD) webserver (https://www.ncbi.nlm.nih.gov/Structure/cdd/wrpsb.cgi) (Marchler-Bauer et al. 2015) with an E-value cut-off set to 10. The trRosetta web server (https://yanglab.nankai.edu.cn/trRosetta/) was used (Xu and Zhang 2012) for *ab initio* modelling. RaptorX and MODELLER were used for secondary structure predictions and structural refinement, respectively. The resulting structures were visualized with the UCSC Chimera 1.15 workflow (Pettersen et al. 2004). RUPEE was used to perform sequence-independent structural comparisons, and HMI-PRED was utilized to infer host-microbe interactions using structural alignment and protein-protein docking methodologies. To compute Mw and isoelectric point (pI), we used the Expasy Compute pI/Mw tool (https://web.expasy.org/compute_pi/) (Gasteiger et al. 2005). N- and O-linked glycosylation were predicted using NetNGlyc (http://www.cbs.dtu.dk/services/NetNGlyc/) (Gupta and Brunak 2001) and NetOGlyc (http://www.cbs.dtu.dk/services/NetOGlyc/) (Steentoft et al. 2013), respectively.

### Translation detection

Assessment of Avo1-9 expression was performed by re-searching LC-MS/MS data from two previously published SARS-CoV-2 studies looking at the infection of human alveolar macrophages (Dalskov et al. 2020) and green monkey Vero E6 cells (Grenga et al. 2020). Data were either search against the human or green monkey SWISS-PROT (Boeckmann et al. 2003) reference proteomes (Human db: 09-2020, 20609 sequences; Green monkey: 08-2020, 19229 sequences) and the UniProt (Consortium 2019) SARS-CoV-2 database (12-2020, 16 sequences) to which the putative Avo1-9 protein sequences had been included. Data were searched using the Mascot search engine (Matrix Science, v.2.5) or by using Proteome Discoverer (Thermo Scientific, v.2.5) employing the Sequest HT and MS Amanda 2.0 search engines. Extended search criteria are included in SM6. Ribosome profiling analyses was performed as described in Ardern *et al.* using data from Finkel et al. (SRR11713356-61, SRR12216748-54).

## Data analysis, visualization, and availability

The bioinformatics workflow, with links to the corresponding software, used in this study can be viewed at https://www.ibp.cz/local/data/avo.

## Acknowledgements

We would like to thank our institution, the University of Ostrava, for an inspiring working environment and academic freedom. Many thanks to Zachary Ardern for help with the Ribo-Seq analysis.

## References

Acevedo JM, Hoermann B, Schlimbach T, Teleman AA. 2018. Changes in global translation elongation or initiation rates shape the proteome via the Kozak sequence. Scientific Reports 8:4018.

Alexandersen S, Chamings A, Bhatta TR. 2020. SARS-CoV-2 genomic and subgenomic RNAs in diagnostic samples are not an indicator of active replication. Nature Communications 11:6059.

Ayoub R, Lee Y. 2019. RUPEE: A fast and accurate purely geometric protein structure search. PLOS One 14:e0213712.

Bahir I, Fromer M, Prat Y, Linial M. 2009. Viral adaptation to host: a proteome-based analysis of codon usage and amino acid preferences. Molecular Systems Biology 5:311.

Balinsky CA, Schmeisser H, Wells AI, Ganesan S, Jin T, Singh K, Zoon KC. 2017. IRAV (FLJ11286), an Interferon-Stimulated Gene with Antiviral Activity against Dengue Virus, Interacts with MOV10. Journal of Virology 91:e01606–16.

Bar-On YM, Flamholz A, Phillips R, Milo R. 2020. SARS-CoV-2 (COVID-19) by the numbers.Eisen MB, editor. eLife 9:e57309.

Beaudoin CA, Jamasb AR, Alsulami AF, Copoiu L, van Tonder AJ, Hala S, Bannerman BP, Thomas SE, Vedithi SC, Torres PHM, et al. 2021. Predicted structural mimicry of spike receptor-binding motifs from highly pathogenic human coronaviruses. Computational and Structural Biotechnology Journal 19:3938–3953.

Boeckmann B, Bairoch A, Apweiler R, Blatter M-C, Estreicher A, Gasteiger E, Martin MJ, Michoud K, O’Donovan C, Phan I. 2003. The SWISS-PROT protein knowledgebase and its supplement TrEMBL in 2003. Nucleic Acids Research 31:365–370.

Bourret J, Alizon S, Bravo IG. 2019. COUSIN (COdon Usage Similarity INdex): A Normalized Measure of Codon Usage Preferences. Genome Biology and Evolution 11:3523–3528.

Chang T-H, Huang H-Y, Hsu JB-K, Weng S-L, Horng J-T, Huang H-D. 2013. An enhanced computational platform for investigating the roles of regulatory RNA and for identifying functional RNA motifs. BMC Bioinformatics 14 Suppl 2:S4.

Consortium U. 2019. UniProt: a worldwide hub of protein knowledge. Nucleic Acids Research 47:D506–D515.

Dalskov L, Møhlenberg M, Thyrsted J, Blay-Cadanet J, Poulsen ET, Folkersen BH, Skaarup SH, Olagnier D, Reinert L, Enghild JJ, et al. 2020. SARS-CoV-2 evades immune detection in alveolar macrophages. EMBO Reports:e51252.

Dilucca M, Forcelloni S, Georgakilas AG, Giansanti A, Pavlopoulou A. 2020. Codon Usage and Phenotypic Divergences of SARS-CoV-2 Genes. Viruses 12:E498.

Dinan AM, Lukhovitskaya NI, Olendraite I, Firth AE. 2020. A case for a negative-strand coding sequence in a group of positive-sense RNA viruses. Virus Evolution 6:veaa007.

Drayman N, Glick Y, Ben-nun-shaul O, Zer H, Zlotnick A, Gerber D, Schueler-Furman O, Oppenheim A. 2013. Pathogens use structural mimicry of native host ligands as a mechanism for host receptor engagement. Cell Host Microbe 14:63–73.

Finkel Y, Mizrahi O, Nachshon A, Weingarten-Gabbay S, Morgenstern D, Yahalom-Ronen Y, Tamir H, Achdout H, Stein D, Israeli O, et al. 2021. The coding capacity of SARS-CoV-2. Nature 589:125–130.

Fuchs U, Rehkamp G, Haas OA, Slany R, Kōnig M, Bojesen S, Bohle RM, Damm-Welk C, Ludwig WD, Harbott J, et al. 2001. The human formin-binding protein 17 (FBP17) interacts with sorting nexin, SNX2, and is an MLL-fusion partner in acute myelogeneous leukemia. Proceedings of the National Academy of Sciences 98:8756–8761.

Gasteiger E, Hoogland C, Gattiker A, Wilkins MR, Appel RD, Bairoch A. 2005. Protein identification and analysis tools on the ExPASy server. In: The proteomics protocols handbook. Springer. p. 571–607.

Gebhardt A, Habjan M, Benda C, Meiler A, Haas DA, Hein MY, Mann A, Mann M, Habermann B, Pichlmair A. 2015. mRNA export through an additional cap-binding complex consisting of NCBP1 and NCBP3. Nature Communications 6:8192.

Goldberg TL, Sibley SD, Pinkerton ME, Dunn CD, Long LJ, White LC, Strom SM. 2019. Multidecade Mortality and a Homolog of Hepatitis C Virus in Bald Eagles (Haliaeetus leucocephalus), the National Bird of the USA. Scientific Reports 9:14953.

Gong Y-N, Chen G-W, Chen C-J, Kuo R-L, Shih S-R. 2014. Computational Analysis and Mapping of Novel Open Reading Frames in Influenza A Viruses. PLOS One 9:e115016.

Gordon DE, Jang GM, Bouhaddou M, Xu J, Obernier K, White KM, O’Meara MJ, Rezelj VV, Guo JZ, Swaney DL, et al. 2020. A SARS-CoV-2 protein interaction map reveals targets for drug repurposing. Nature 583:459–468.

Gouveia D, Grenga L, Gaillard J-C, Gallais F, Bellanger L, Pible O, Armengaud J. 2020. Shortlisting SARS-CoV-2 Peptides for Targeted Studies from Experimental Data-Dependent Acquisition Tandem Mass Spectrometry Data. PROTEOMICS 20:2000107.

Gregersen LH, Schueler M, Munschauer M, Mastrobuoni G, Chen W, Kempa S, Dieterich C, Landthaler M. 2014. MOV10 Is a 5′ to 3′ RNA Helicase Contributing to UPF1 mRNA Target Degradation by Translocation along 3′ UTRs. Molecular Cell 54:573–585.

Grenga L, Gallais F, Pible O, Gaillard J-C, Gouveia D, Batina H, Bazaline N, Ruat S, Culotta K, Miotello G, et al. 2020. Shotgun proteomics analysis of SARS-CoV-2-infected cells and how it can optimize whole viral particle antigen production for vaccines. Emerging Microbes & Infections 9:1712–1721.

Gu H, Chu DKW, Peiris M, Poon LLM. 2020. Multivariate analyses of codon usage of SARS-CoV-2 and other betacoronaviruses. Virus Evolution 6:veaa032.

Guarracino A, Pepe G, Ballesio F, Adinolfi M, Pietrosanto M, Sangiovanni E, Vitale I, Ausiello G, Helmer-Citterich M. 2021. BRIO: a web server for RNA sequence and structure motif scan. Nucleic Acids Research 49:W67–W71.

Gupta R, Brunak S. 2001. Prediction of glycosylation across the human proteome and the correlation to protein function. In: Pac Symp Biocomput. Vol. 7. p. 310–22.

Guven-Maiorov E, Hakouz A, Valjevac S, Keskin O, Tsai C-J, Gursoy A, Nussinov R. 2020. HMI-PRED: a web server for structural prediction of host-microbe interactions based on interface mimicry. Journal of Molecular Biology 432:3395–3403.

Hickman HD, Mays JW, Gibbs J, Kosik I, Magadán JG, Takeda K, Das S, Reynoso GV, Ngudiankama BF, Wei J, et al. 2018. Influenza A Virus Negative Strand RNA Is Translated for CD8+ T Cell Immunosurveillance. The Journal of Immunology 201:1222–1228.

Hu B, Guo H, Zhou P, Shi Z-L. 2020. Characteristics of SARS-CoV-2 and COVID-19. Nature Reviews Microbiology:1–14.

Irigoyen N, Firth AE, Jones JD, Chung BY-W, Siddell SG, Brierley I. 2016. High-Resolution Analysis of Coronavirus Gene Expression by RNA Sequencing and Ribosome Profiling. PLOS Pathogens 12:e1005473.

Jaafar ZA, Kieft JS. 2019. Viral RNA structure-based strategies to manipulate translation. Nature Reviews Microbiology 17:110–123.

Jiang H, Li Y, Zhang H, Wang W, Yang X, Qi H, Li H, Men D, Zhou J, Tao S. 2020. SARS-CoV-2 proteome microarray for global profiling of COVID-19 specific IgG and IgM responses. Nature Communications 11:1–11.

Jitobaom K, Phakaratsakul S, Sirihongthong T, Chotewutmontri S, Suriyaphol P, Suptawiwat O, Auewarakul P. 2020. Codon usage similarity between viral and some host genes suggests a codon-specific translational regulation. Heliyon 6:e03915.

Källberg M, Wang H, Wang S, Peng J, Wang Z, Lu H, Xu J. 2012. Template-based protein structure modeling using the RaptorX web server. Nature protocols 7:1511–1522.

Komazin-Meredith G, Santos WL, Filman DJ, Hogle JM, Verdine GL, Coen DM. 2008. The Positively Charged Surface of Herpes Simplex Virus UL42 Mediates DNA Binding *. Journal of Biological Chemistry 283:6154–6161.

Krogh A, Larsson B, von Heijne G, Sonnhammer EL. 2001. Predicting transmembrane protein topology with a hidden Markov model: application to complete genomes. Journal of Molecular Biology 305:567–580.

La Bella T, Imbeaud S, Peneau C, Mami I, Datta S, Bayard Q, Caruso S, Hirsch TZ, Calderaro J, Morcrette G, et al. 2020. Adeno-associated virus in the liver: natural history and consequences in tumour development. Gut 69:737–747.

Li Y, Masaki T, Shimakami T, Lemon SM. 2014. hnRNP L and NF90 interact with hepatitis C virus 5′-terminal untranslated RNA and promote efficient replication. Journal of Virology 88:7199–7209.

Li Y, Yang Xinai, Wang N, Wang H, Yin B, Yang Xiaoping, Jiang W. 2020. GC usage of SARS-CoV-2 genes might adapt to the environment of human lung expressed genes. Molecular Genetics and Genomics: MGG 295:1537–1546.

Lomize MA, Pogozheva ID, Joo H, Mosberg HI, Lomize AL. 2012. OPM database and PPM web server: resources for positioning of proteins in membranes. Nucleic Acids Research 40:D370–376.

Marchler-Bauer A, Derbyshire MK, Gonzales NR, Lu S, Chitsaz F, Geer LY, Geer RC, He J, Gwadz M, Hurwitz DI. 2015. CDD: NCBI’s conserved domain database. Nucleic Acids Research 43:D222–D226.

May JP, Simon AE. 2021. Targeting of viral RNAs by Upf1-mediated RNA decay pathways. Current Opinion in Virology 47:1–8.

Miyazono K, Zhi Y, Takamura Y, Nagata K, Saigo K, Kojima T, Tanokura M. 2010. Cooperative DNA-binding and sequence-recognition mechanism of aristaless and clawless. The EMBO journal 29:1613–1623.

Monjaret F, Bourg N, Suel L, Roudaut C, Le Roy F, Richard I, Charton K. 2014. Cis-splicing and translation of the pre-trans-splicing molecule combine with efficiency in spliceosome-mediated RNA trans-splicing. Molecular Therapy 22:1176–1187.

Nair VP, Anang S, Subramani C, Madhvi A, Bakshi K, Srivastava A, Nayak B, CT RK, Surjit M. 2016. Endoplasmic reticulum stress induced synthesis of a novel viral factor mediates efficient replication of genotype-1 hepatitis E virus. PLOS Pathogens 12:e1005521.

Narita R, Takahasi K, Murakami E, Hirano E, Yamamoto SP, Yoneyama M, Kato H, Fujita T. 2014. A Novel Function of Human Pumilio Proteins in Cytoplasmic Sensing of Viral Infection. PLOS Pathogens 10:e1004417.

Nelson CW, Ardern Z, Goldberg TL, Meng C, Kuo C-H, Ludwig C, Kolokotronis S-O, Wei X. 2020. Dynamically evolving novel overlapping gene as a factor in the SARS-CoV-2 pandemic.Rokas A, Wittkopp PJ, Piontkivska H, editors. eLife 9:e59633.

Noderer WL, Flockhart RJ, Bhaduri A, Diaz de Arce AJ, Zhang J, Khavari PA, Wang CL. 2014. Quantitative analysis of mammalian translation initiation sites by FACS-seq. Molecular Systems Biology 10:748.

Park S-J, Lee Jumin, Qi Y, Kern NR, Lee HS, Jo S, Joung I, Joo K, Lee Jooyoung, Im W. 2019. CHARMM-GUI Glycan Modeler for modeling and simulation of carbohydrates and glycoconjugates. Glycobiology 29:320–331.

Pettersen EF, Goddard TD, Huang CC, Couch GS, Greenblatt DM, Meng EC, Ferrin TE. 2004. UCSF Chimera—a visualization system for exploratory research and analysis. Journal of Computational Chemistry 25:1605–1612.

Puigbò P, Bravo IG, Garcia-Vallve S. 2008. CAIcal: A combined set of tools to assess codon usage adaptation. Biology Direct 3:38.

Reily C, Stewart TJ, Renfrow MB, Novak J. 2019. Glycosylation in health and disease. Nature Reviews Nephrology 15:346–366.

Renuse S, Vanderboom PM, Maus AD, Kemp JV, Gurtner KM, Madugundu AK, Chavan S, Peterson JA, Madden BJ, Mangalaparthi KK, et al. 2021. A mass spectrometry-based targeted assay for detection of SARS-CoV-2 antigen from clinical specimens. EBioMedicine 69:103465.

Requião RD, Carneiro RL, Moreira MH, Ribeiro-Alves M, Rossetto S, Palhano FL, Domitrovic T. 2020. Viruses with different genome types adopt a similar strategy to pack nucleic acids based on positively charged protein domains. Scientific Reports 10:5470.

Roy A, Guo F, Singh B, Gupta S, Paul K, Chen X, Sharma NR, Jaishee N, Irwin DM, Shen Y. 2021. Base Composition and Host Adaptation of the SARS-CoV-2: Insight From the Codon Usage Perspective. Frontiers in Microbiology 12:548275.

Salamov AA, Nishikawa T, Swindells MB. 1998. Assessing protein coding region integrity in cDNA sequencing projects. Bioinformatics 14:384–390.

Sali A, Blundell TL. 1993. Comparative protein modelling by satisfaction of spatial restraints. Journal of Molecular Biology 234:779–815.

Sharp PM, Li WH. 1987. The codon Adaptation Index--a measure of directional synonymous codon usage bias, and its potential applications. Nucleic Acids Research 15:1281–1295.

Shi M, Jagger BW, Wise HM, Digard P, Holmes EC, Taubenberger JK. 2012. Evolutionary Conservation of the PA-X Open Reading Frame in Segment 3 of Influenza A Virus. Journal of Virology 86:12411–12413.

Steentoft C, Vakhrushev SY, Joshi HJ, Kong Y, Vester-Christensen MB, Schjoldager KT-B, Lavrsen K, Dabelsteen S, Pedersen NB, Marcos-Silva L. 2013. Precision mapping of the human O-GalNAc glycoproteome through SimpleCell technology. The EMBO journal 32:1478–1488.

Szczesny RJ, Borowski LS, Brzezniak LK, Dmochowska A, Gewartowski K, Bartnik E, Stepien PP. 2010. Human mitochondrial RNA turnover caught in flagranti: involvement of hSuv3p helicase in RNA surveillance. Nucleic Acids Research 38:279–298.

Tan K-E, Ng WL, Marinov GK, Yu KH-O, Tan LP, Liau ES, Goh SY, Yeo KS, Yip KY, Lo K-W, et al. 2021. Identification and characterization of a novel Epstein-Barr Virus-encoded circular RNA from LMP-2 Gene. Scientific Reports 11:14392.

Thomas C, Moraga I, Levin D, Krutzik PO, Podoplelova Y, Trejo A, Lee C, Yarden G, Vleck SE, Glenn JS, et al. 2011. Structural linkage between ligand discrimination and receptor activation by type I interferons. Cell 146:621–632.

Veltri D, Wight MM, Crouch JA. 2016. SimpleSynteny: a web-based tool for visualization of microsynteny across multiple species. Nucleic Acids Research 44:W41–W45.

Webb B, Sali A. 2017. Protein structure modeling with MODELLER. In: Functional genomics. Springer. p. 39–54.

Wu F, Zhao S, Yu B, Chen Y-M, Wang W, Song Z-G, Hu Y, Tao Z-W, Tian J-H, Pei Y-Y. 2020. A new coronavirus associated with human respiratory disease in China. Nature 579:265–269.

Xu D, Zhang Y. 2012. Ab initio protein structure assembly using continuous structure fragments and optimized knowledge-based force field. Proteins: Structure, Function, and Bioinformatics 80:1715–1735.

Yang J, Anishchenko I, Park H, Peng Z, Ovchinnikov S, Baker D. 2020. Improved protein structure prediction using predicted interresidue orientations. Proceedings of the National Academy of Sciences 117:1496–1503.

Zhang J, Cruz-cosme R, Zhuang M-W, Liu D, Liu Y, Teng S, Wang P-H, Tang Q. 2020. A systemic and molecular study of subcellular localization of SARS-CoV-2 proteins. Signal Transduction and Targeted Therapy 5:1–3.

Zhang Y, Skolnick J. 2005. TM-align: a protein structure alignment algorithm based on the TM-score. Nucleic Acids Research 33:2302–2309.

Zheng J. 2020. SARS-CoV-2: an emerging coronavirus that causes a global threat. International Journal of Biological Sciences 16:1678.

Zuallaert J, Kim M, Soete A, Saeys Y, Neve WD. 2018. TISRover: ConvNets learn biologically relevant features for effective translation initiation site prediction. International Journal of Data Mining and Bioinformatics 20:267–284.

